# Dorsal Anterior Cingulate Cortex Encodes the Integrated Incentive Motivational Value of Cognitive Task Performance

**DOI:** 10.1101/2020.09.20.305482

**Authors:** Debbie M. Yee, Jennifer L. Crawford, Bidhan Lamichhane, Todd S. Braver

**Author notes:** Correspondence should be addressed to: Debbie Yee, Brown University, Cognitive, Linguistic, and Psychological Sciences, 190 Thayer Street, Providence, RI, 02912. **Author Contributions**: D.M.Y. and T.S.B. designed research; D.M.Y. and J.L.C. performed research; D.M.Y. and J.L.C., and B.L. analyzed data; D.M.Y. wrote first draft of the manuscript; D.M.Y., J.L.C., B.L., and T.S.B. edited manuscript.

## Abstract

Humans can seamlessly combine value signals from diverse motivational incentives, yet it is not well-understood how these signals are “bundled” in the brain to modulate cognitive control. The dorsal anterior cingulate cortex (dACC) is theorized to integrate motivational value dimensions in the service of goal-directed action, though this hypothesis has yet to receive rigorous confirmation. In the present study, we examined the role of human dACC in motivational incentive integration. Healthy young adult men and women were scanned with fMRI while engaged in an experimental paradigm that quantifies the combined effects of liquid (e.g., juice, neutral, saltwater) and monetary incentives on cognitive task performance. Monetary incentives modulated trial-by-trial dACC activation, whereas block-related effects of liquid incentives on dACC activity were observed. When bundled together, incentive-related dACC modulation predicted fluctuations in both cognitive performance and self-report motivation ratings. Statistical mediation analyses suggest that dACC encoded the incentives in terms of their integrated subjective motivational value, and that this value signal was most proximally associated with task performance. Finally, we confirmed that these incentive integration effects were selectively present in dACC. Together, the results support an account in which dACC integrates motivational signals to compute the expected value of goal-directed cognitive control.

**Significance Statement:** How are primary and secondary incentives integrated in the brain to influence goal-directed behavior? Using an innovative experimental fMRI paradigm that combines motivational incentives that have historically been studied independently between species (e.g., monetary rewards for humans, food rewards for animals), we examine the relationship between incentive motivational value and cognitive control allocation. We find evidence that the integrated incentive motivational value of combined incentives is encoded in human dorsal anterior cingulate cortex (dACC). Further, self-reported motivational shifts mediated the effects of incentive-modulated dACC activity on task performance, revealing convergence in how self-reported and experimentally-induced motivation are encoded in the human brain. Our findings may inform future translational studies examining affective/motivational and cognitive impairments in psychopathology (e.g., anxiety, depression, addiction).

## Introduction

A remarkable aspect of the interaction between motivation and cognition is the seamless ability that humans have in integrating diverse motivational incentives when allocating cognitive resources towards pursuit of mentally demanding goals (Botvinick & Braver, 2015). For example, when working towards a challenging project, a worker might be motivated by a potential raise, the praise received from their supervisor, the tasty snack they promised themselves upon completion, or most likely, a combination of all three. Although individuals likely “bundle” values from multiple incentives to influence goal pursuit (FitzGerald et al., 2009), prior studies of human motivation have primarily examined monetary rewards (Bahlmann et al., 2015; Kouneiher et al., 2009; Padmala & Pessoa, 2011). Few studies have explored how biological incentives (e.g., food/drink) influence cognitive task performance (Krug & Braver, 2014). Yet, consideration of primary and “bundled” incentives is theoretically important for clarifying how incentive motivational value is encoded in the brain how it can modulate goal pursuit. In this human fMRI study, we investigate the neural mechanisms underlying effects of integrated motivational incentives on cognitive control.

Several theoretical frameworks provide relevant predictions for neural mechanisms underpinning incentive integration. In neuroeconomics, the “common currency” account suggests diverse incentives are represented in a common neural representation that enables incentives to be compared, combined, and selected under decision-making contexts (Levy & Glimcher, 2012; Padoa-Schioppa & Cai, 2011). The explicit encoding of incentives may reflect a valuation process that enables rank ordering based upon subjective utility. One natural form for this utility encoding might be subjective motivational value, which is putatively represented in ventromedial prefrontal cortex and striatum (Chib et al., 2009; Sescousse et al., 2015). The encoding of incentives as subjective motivational value is relevant for theories of motivation-cognition interaction, which suggests cognitive control is tightly coupled with motivational signals (Parro et al., 2018; Yee & Braver, 2018). Conversely, the Expected Value of Control (EVC) theory postulates dorsal anterior cingulate cortex (dACC) plays a key role in integrating positive and negative outcomes to modulate cognitive control signals (Shenhav et al., 2013, 2017). Previous studies have demonstrated that dACC is engaged during incentivized cognitive tasks (Parro et al., 2018) and motivated action selection (Holroyd & Yeung, 2012; Rushworth et al., 2004). Further, dACC is sensitive to reward and punishment (Fujiwara et al., 2009; Lake et al., 2019), and to benefits and costs associated with cognitive control (Sayalı & Badre, 2018; Westbrook et al., 2019). Nevertheless, to robustly test whether dACC is an integrative motivation-cognition hub, it is necessary to demonstrate dACC integrates the value of diverse incentives with both positive and negative motivational value, and that dACC signals are associated with fluctuations in self-reported motivation and cognitive task performance. A rigorous formal investigation of incentive integration requires an experimental paradigm that parametrically bundles incentives to measure their modulation of cognitive task performance.

We recently developed a novel incentive integration paradigm which has these critical components, enabling a rigorous test of whether dACC serves as a motivation-cognitive control hub. In this paradigm, liquid and monetary incentives are “bundled” together, but independently manipulated in a trial-by-trial and block-wise manner, enabling estimation of incentive-modulated effects on both self-reported motivation and cognitive task performance. The incorporation of liquids enables straightforward examination of motivational valence effects through utilization of positive/appetitive (juice), neutral (tasteless liquid), and negative/aversive (saltwater) incentives. Since participants directly consume the liquids, their subjective ratings of these liquids may indicate their motivational influence on task performance. Prior work has demonstrated that both incentives are additively combined to influence self-reported motivation and task performance (Yee et al., 2016, 2019). We leverage this paradigm in a human fMRI study with healthy young participants to test the compatibility of dACC activity patterns a key claim of the EVC account: that dACC integrates potential positive and negative values across diverse incentives to adjust the control signal; that is, as integrated subjective motivational value increases, so should dACC activity and the subsequent allocation of control. To preview, our results are consistent with this prediction, suggesting that dACC modulates motivation-cognition interactions via internal representation of bundled motivational value signals, and this representation is what enables motivation-linked modulation of task performance under high cognitive control demands.

## Materials and Methods

### Experimental Design

#### Participants

51 right-handed participants (25 female; 18-38 years, M=25.1, SD=4.8) with normal or corrected-to-normal vision participated in the experiment. All participants provided written consent approved by the Washington University Institutional Review Board, and received payment for their participation ($25 per hour), plus additional earnings of up to 15 dollars based on task performance. Five participants were excluded from analyses due to technical error, participant inability to complete the task, or participant noncompliance with task instructions. The final sample consisted of 46 participants (22 females; 18–38 years, M=25.4, SD=4.9). All demographic and self-report data were collected and managed using a secure web-based application, Research Electronic Data Capture (REDCap), hosted at Washington University (Harris et al., 2009).

#### Incentive Integration Task

To examine the dissociable and bundled effects of primary and secondary incentives on cognitive control, we adapted the consonant-vowel odd-even (CVOE) cued task-switching paradigm developed by Yee et al. (2016, 2019). On each trial, a letter-number pair was visually presented (e.g., one letter and one number on the screen), and participants were tasked with categorizing the target symbol based on the task instruction briefly presented at the beginning of each trial (e.g., classify the letter was a vowel or consonant, the number as odd or even). The task for a given trial was indicated by a cue display, which was randomized across trials and preceded the number-letter pair, indicating either “Attend Letter” or “Attend Number”. Participants maintained the current task and associated response rules in working memory during a subsequent blank cue-target interval. A monetary reward cue was also presented each trial, placed above and below the task cue, which indicated whether the trial was associated with a low, medium, or high reward value (displayed as “$”, “$$”, or “$$$$”). The values of the monetary reward cues were randomized across trials. Although reward cues were always presented with task cues, participants could only earn monetary reward during incentive blocks (i.e., not during practice and baseline blocks).

During the incentive blocks, participants could earn monetary rewards for fast and accurate task performance (See Procedure). One key aspect of the experimental design was the utilization of monetary reward cues that varied on a trial-by-trial basis. A second key aspect was that successful attainment of the monetary reward was indicated by oral liquid delivery to the participant’s mouth as post-trial performance feedback. At the end of trials in which participants were accurate and faster than the criterion RT, they received a 1 mL drop of liquid directly to their mouths. Participants only received liquid feedback for successfully earning monetary reward in a given trial, and did not receive liquid if they were incorrect, too slow, or did not respond. Importantly, although the type of liquid received was blocked, such that the liquid feedback could be positive/appetitive (apple juice), neutral (isotonic tasteless solution), or negative/aversive (saltwater), the symbolic meaning of the liquid was kept constant. Thus, any behavioral differences observed between liquid types can be attributed to differential subjective valuation of liquid feedback, and simultaneous consideration of both monetary rewards and liquid incentives during task performance. Thus, because receipt of both monetary reward and liquid feedback was performance-contingent, participants integrated the value of both incentives (i.e., motivational incentive integration) when performing the task. Thus, the task enables straightforward comparison of the parametric effects of value on task performance for each motivational incentive (e.g., low vs. medium vs. high monetary rewards), as well as for “bundled” incentives (e.g., juice + high monetary reward vs. neutral + high monetary reward) that reflect the effect of integrated motivational value on cognitive task performance.

The task was programmed with Psychtoolbox 3 (version 3.0.12) in Matlab (version 2016b) and displayed on the projector connected to a laptop computer. Each trial consisted of a fixation cross displayed for 300 ms, a cue display with the task instruction and monetary reward value for 500 ms, a blank display for 4000 ms (cue-target interval), a target display of the number-letter pair for 2000 ms, a second fixation display for 1000 ms, and a feedback display for 2000 ms (See Figure 1A). Finally, an inter-trial interval of randomized duration (3000, 5000, or 7000 ms) displayed the fixation cross prior to the start of the next trial. Response mappings were counterbalanced between participants. A more detailed schematic of the task trial with all of the timing variables is included in the extended data (See Figure 1-1).

**Figure 1:**
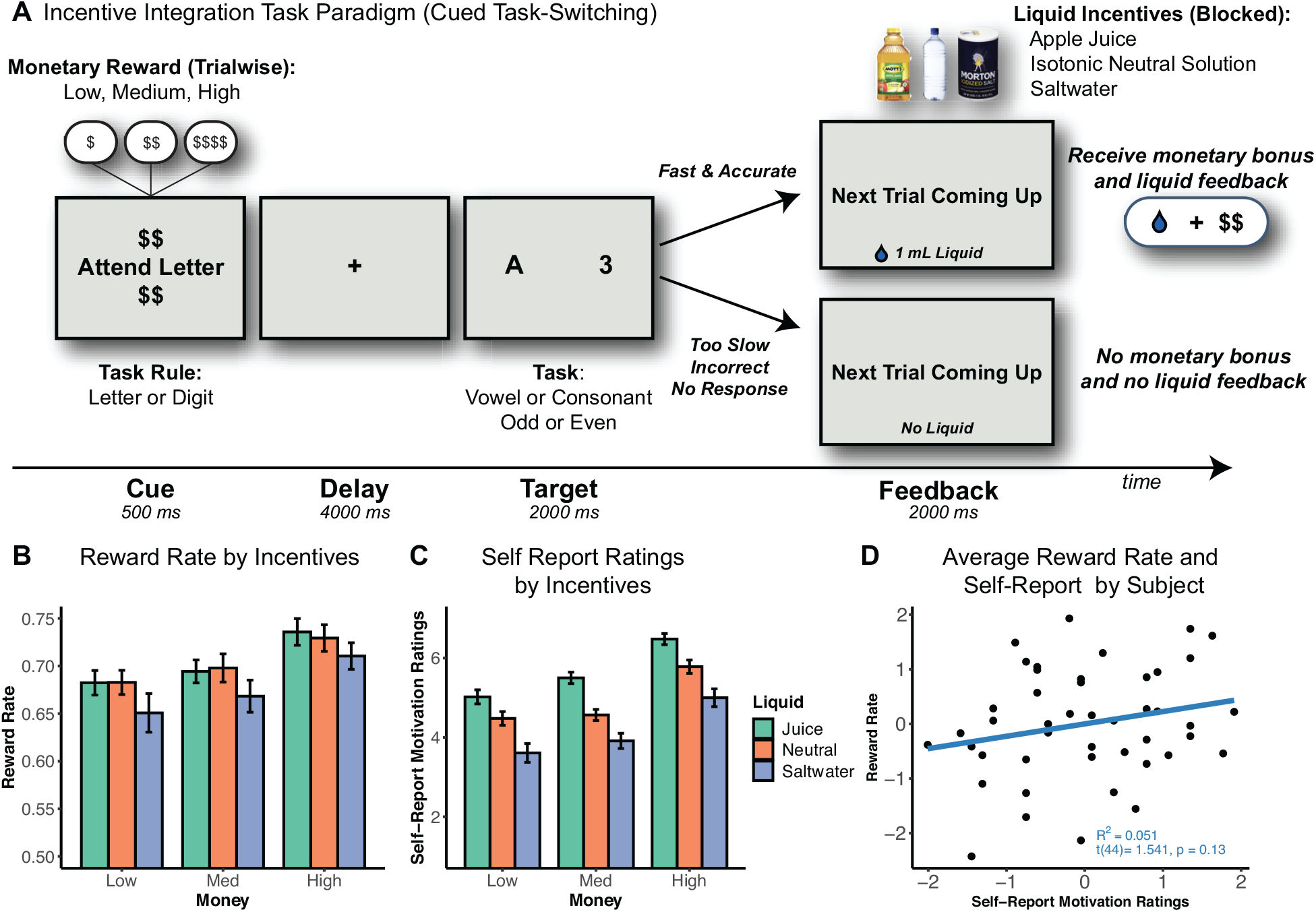
Incentive Integration Task Paradigm and Behavioral Results. **A)** Incentive integration task paradigm. Participants performed letter-digit cued task-switching and could earn monetary rewards and liquid incentives for accurate and fast performance (below an individualized criterion threshold). Notably, as the receipt of both monetary reward and liquid feedback was performance-contingent, participants had to integrate the value of both types of motivational incentives when performing this cognitive task. **B)** Reward rate by motivational incentive conditions. Participants performed better with trials with higher monetary reward. In terms of liquid effects, participants performed worse on saltwater compared to juice or neutral trials. Error bars indicate SEM. **C)** Self-Report Motivation Ratings by motivational incentive conditions. Participants reported they were more motivated for higher monetary reward and more appetitive liquid incentives. Error bars indicate SEM. A hierarchical regression revealed that the inclusion of motivation ratings significantly predicted variance in reward rate beyond the experimental effects. **D)** Scatterplot of z-scored averaged reward rate and self-report motivation ratings by participant. Notably, although both measures are sensitive to the motivational incentive conditions, there was only a weak positive association between self-reported motivation and reward rate. These data suggest that the two measures reflect overlapping yet dissociable motivational components of the incentivized cognitive task.

**Figure 1-1:**
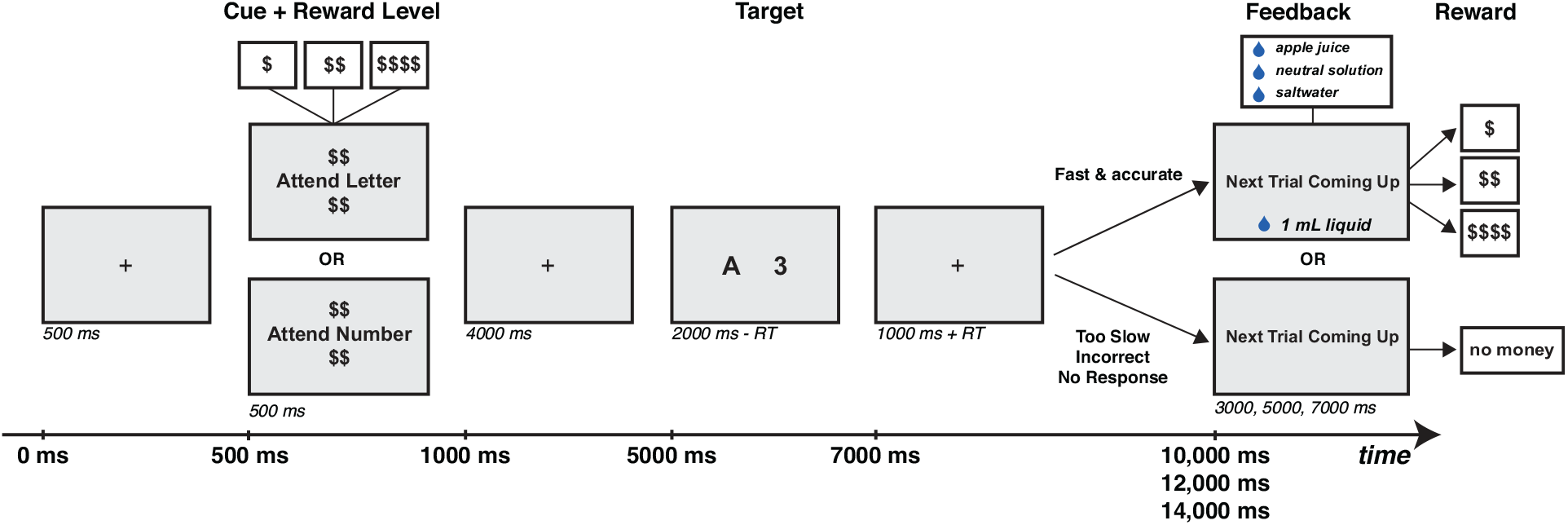
Detailed Schematic of Single Trial from Incentive Integration Task. Adapted for the MRI scanner. Participants performed a consonant-vowel odd-even (CVOE) switching task, which entailed being presented with an ambiguous letter-number pair, and being asked to categorize the target symbol based on the task cue preceding the target (e.g., “Attend Number” or “Attend Letter). A reward cue was placed above and below each instruction cue, which indicated low ($), medium ($$), or high ($$$$) reward. Monetary reward cues were randomized across trials within each block. Importantly, the dollar signs associated with the cues indicated how much the subject could earn on each trial. If subjects were accurate and faster than a subject criterion response time (40% of fastest correct response times for all trials during the baseline block), then they received 1 mL of liquid as performance feedback at the end of the trial. If subjects answered incorrectly, too slowly, or not at all, they neither received monetary reward or liquid. Liquid type was manipulated in a blocked fashion, counterbalanced across subjects, and was positive (apple juice), neutral (isotonic tasteless solution), or negative (saltwater). Because the liquid delivery is symbolic, which means that it conveys the same information regardless of the type, any differences across liquids would suggest that subjects are incorporating the value of that liquid to influence their task performance. Thus, this task design enables us to test for the dissociable and integrated effects of monetary and liquid incentives on cognitive task performance.

#### Procedure

Participants were asked to abstain from eating or drinking anything besides water for two hours before the start of the session. Upon arrival, participants completed a contact information questionnaire with demographic information, along with the Behavioral Inhibition & Avoidance Scales (BIS/BAS), a self-report survey often used to measure individual differences in motivation to avoid aversive outcomes and approach goal-oriented outcomes (Carver & White, 1994).

Next, participants practiced the task for 30 minutes in a testing room. They first practiced single tasks (letter categorization or number categorization only; order counterbalanced across participants), followed by practice of a mixed task block (task-switching between letter and number task rules). During the three practice blocks (one letter, one number, one mixed), participants received visual performance feedback to indicate accurate performance after each trial. During the three baseline blocks (one letter, one number, one mixed), participants no longer received performance feedback. Additionally, participants practiced swallowing the liquids while lying down on a bed in the testing room, and several drops of the neutral solution liquid were delivered via a pacifier and plastic tubing from the computer-triggered liquid delivery setup.

During scanning, a mixed block/event-related task design was used to optimize characterization of nonlinear and time-sensitive neuronal responses, and to enable simultaneous extraction of trial-related transient activity and block-related sustained activity related to task-level processing (Petersen & Dubis, 2012). Each run consisted of 48 trials divided between three blocks (16 trials per block), alternating with 30 seconds of “rest” in which participants were instructed to attend a fixation cross display (See Figure 2B). Participants performed one or two baseline runs, in which they performed the task without earning incentives. Each participant’s reward criterion (40%) was calculated based on their response times in the baseline runs. Following the baseline runs, participants completed six incentive blocks with liquid delivered as performance feedback for the successful attainment of monetary reward (two juice, two neutral, two saltwater). Liquid order was counterbalanced. Participants received 1 mL of liquid delivered directly to their mouths (see Liquid Setup and Delivery Procedure) if they were accurate and faster than their calculated reward criterion (40% faster than baseline performance). If participants were incorrect, too slow, or did not respond, they did not earn monetary reward and did not receive any liquid during that trial.

**Figure 2:**
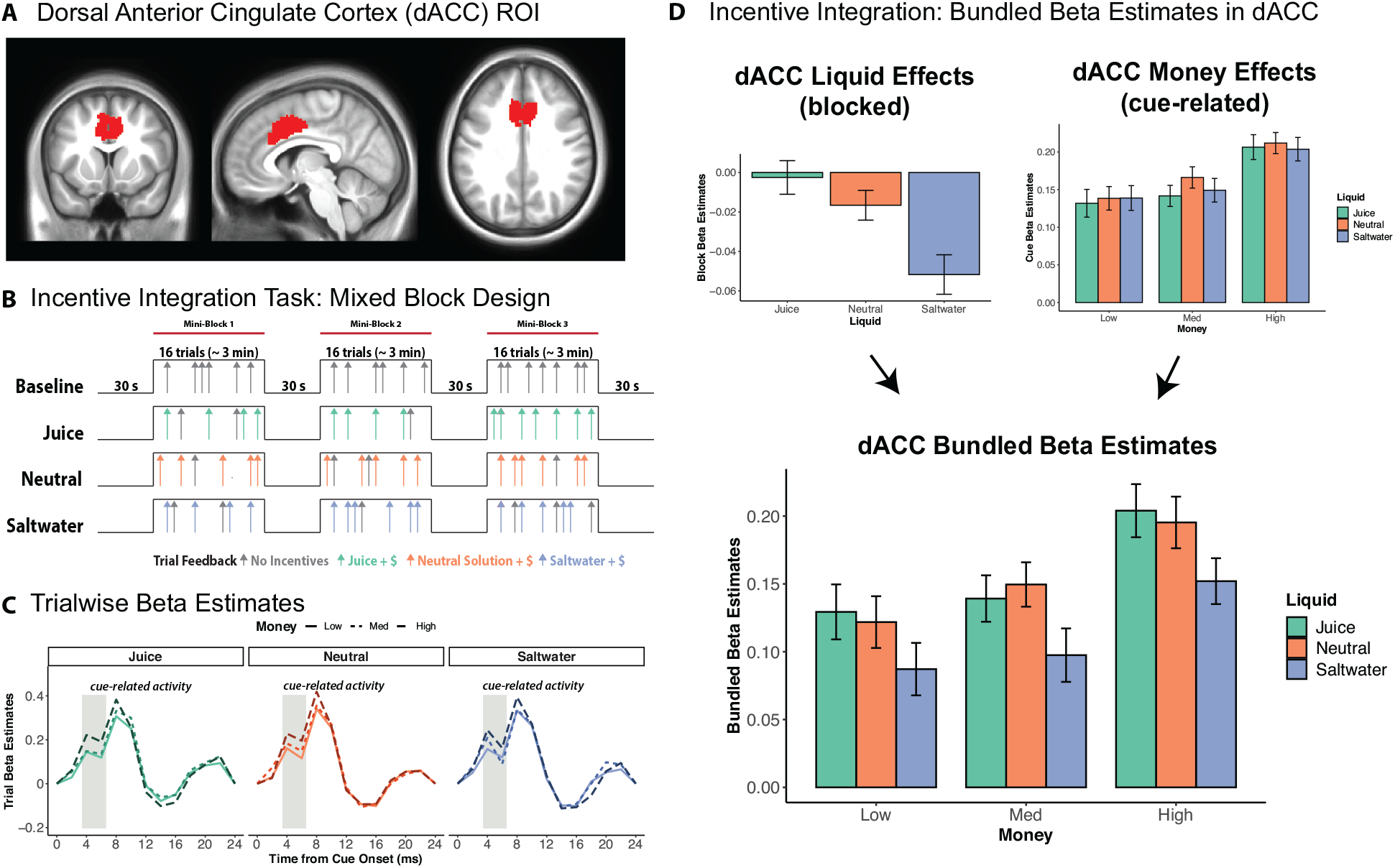
Dorsal ACC Encodes Both Monetary Rewards and Liquid Incentives in Bundled Beta Estimates. **A)** Bilateral dACC ROI mask. This ROI encompasses the peak voxels of dACC based upon a prior meta-analysis on motivated cognitive control (Parro, Dixon and Christoff, 2018). **B)** Mixed block/event-related design for incentive integration task. Participants performed 8 task blocks total, with two baseline blocks and six incentive blocks. Each participant’s reward criterion (40%) was calculated based upon performance in the baseline run and used to determine the RT threshold by which fast and accurate performance would lead to earned monetary and liquid incentives. Monetary reward value randomly varied on a trialwise basis. Liquid type was blocked and counterbalanced and delivered as performance feedback for successful attainment of monetary reward (shown in colored arrows). Participants did not receive money nor liquid for slow, incorrect, or abstained responses. A GLM was applied to extract the beta estimates for the sustained liquid block conditions and transient event-related motivation conditions for each participant (Petersen and Dubis, 2012). **C)** Trial wise beta estimates are illustrated for each of the 9 motivational incentive conditions. Darker colors indicate higher monetary reward level. Cue-related activity is highlighted in gray rectangles in each plot, demonstrating a significant monetary reward effect 4-6 seconds after cue onset. **D)** Bundled betas estimates in dACC are calculated by combining the beta estimates for sustained liquid effects and cue-related monetary reward effects from the event-related hemodynamic response. We assume an additive relationship between the monetary and liquid effects in terms of BOLD signal representation of integrated incentive value. Reward rate was significantly predicted by dACC bundled betas, revealing that dACC represented the aggregate motivational value of primary and secondary incentives and is associated with parametric modulation of motivated cognitive task performance.

After completion of the scanning session, participants completed a post-task questionnaire and reported Likert ratings (1-7) of motivation of the liquids for each of the nine task conditions (See Figure 1C). Specifically, participants were asked how motivated they were on each of the 9 motivational incentive conditions (e.g., “How motivated were you on the Juice $ trials?”). Participants additionally reported Likert ratings of liking and intensity for each liquid (See Figure 1-3). Specifically, participants were asked “Please indicate on a scale of 1-7 how much you like or dislike this liquid” and “Please indicate on a scale of 1-7 how intense you find the taste of this liquid.” All self-report questions posed to participants are listed in Table 1-1. Reward earnings were calculated and added to their base rate earnings for experimental participation.

**Table 1-1:** Self-report questions posed at the end of the study. For all the questions, participants completed Likert ratings on a scale from 1 (lowest) to 7 (highest). For the motivation questions, there were 9 questions – one for each of the motivational incentive conditions. For the liking and intensity questions, there were 3 questions – one for each liquid type. For each liquid, participants were asked to rate how much the liked the liquid from 1 (least liked) to 7 (most liked), as well as how intense they found the taste of each liquid from 1 (least intense) to 7 (most intense).

|  | Questions |
| --- | --- |
| ***Motivation*** | How motivated were you on *Juice \$* trials?<br>How motivated were you on *Juice \$\$* trials?<br>How motivated were you on *Juice \$\$\$\$* trials?<br>How motivated were you on *Neutral \$* trials?<br>How motivated were you on *Neutral \$\$* trials?<br>How motivated were you on *Neutral \$\$\$\$* trials?<br>How motivated were you on *Saltwater \$* trials?<br>How motivated were you on *Saltwater \$\$* trials?<br>How motivated were you on *Saltwater \$\$\$\$* trials? |
| ***Liking*** | Please indicate on a scale of 1-7 how much you like or dislike this liquid. |
| ***Intensity*** | Please indicate on a scale of 1-7 how intense you find the taste of this liquid. |

#### Liquid Setup and Delivery Procedure

Prior to the scan, the neutral and saltwater liquid solutions were prepared in a testing room in the CCP lab. The isotonic neutral solution consisted of 1 liter of distilled water, .0495g of NaHCO3 (sodium bicarbonate), and .4668g of KCl (potassium chloride). The saltwater solution consisted of 500 mL of distilled water and 8.8g of non-iodized salt. The juice used was 100% apple juice (Mott’s) and purchased from the store.

During both the behavioral practice and scanning session, the liquids were dispensed using a digital infusion pump (model SP210iw, World Precision Instruments, Inc.) triggered by an output signal from the Matlab script and delivered via Tygon tubing and a pacifier directly to the participant’s mouth. The infusion pump was located in the control room, and lengthy tubing was used to ensure that the dispensed liquid was delivered to the participants. The type of liquid delivered was manipulated in a blocked fashion, counterbalanced across participants, such that on a given block participants would receive positive / appetitive (apple juice), neutral (isotonic tasteless solution), or negative / aversive (saltwater) performance feedback.

### Statistical Analysis

#### Behavioral Data Analysis

Behavioral data were analyzed in Rstudio using the R statistical language (RCoreTeam, 2017; RStudioTeam, 2016). All of the data were visualized using the ggplot2 package (Wickham, 2016). All of the linear mixed-effects models were conducted using the lmerTest (Kuznetsova et al., 2015) and LME4 (Bates et al., 2015) packages. We utilized linear mixed models for our behavioral and ROI based analyses to facilitate more straightforward comparisons with analyses conducted in our prior work with this task (Yee et al., 2016, 2019). When applicable, 95% confidence intervals for the effects were calculated using the “confint” function in the lme4 package. The knitr and Rmarkdown packages were used to create dynamic reports of the results (Xie, 2019). The behavioral data and analysis scripts are available on OSF: https://osf.io/upka4/.

#### Bayesian Multilevel Mediation Analysis

We conducted a Bayesian multilevel mediation analyses (BMLM) to test the within-subjects mediated effects between dACC, self-reported motivation ratings, and motivated task performance. We adopted a Bayesian approach for the multilevel mediation analysis as it allows for more precise estimates of indirect effects, as the indirect effects typically do not follow a normal sampling distribution. Moreover, BMLM analyses are more conceptually straightforward and allow for simulation of sampling distributions to estimate parameters and credible intervals that characterize the mediated effects between our variable of interest (Yuan & MacKinnon, 2009). All variables were within-person centered to extract within-subject deviations across the nine motivational conditions, which captured fluctuations relative to each participant’s subject means. The Bayesian multilevel mediation analyses were conducted in R using the bmlm package in R (Vuorre & Bolger, 2017), which depends on the powerful Stan Bayesian inference engine. We used the default priors from the bmlm package (i.e., default priors for regression coefficients were normally distributed (*Normal*(0,1000)) and group-level SDs were Cauchy distributed (*Cauchy*(0,50)). The models were implemented with 10,000 samples drawn from the posterior distribution for each of the 4 MCMC chains. Half of the samples (5,000) were used for sampling (which is the default). All Rhat values were equal to 1.00, indicating accurate estimates of the posterior distribution and model convergence. We report the medians and the 90% credible intervals from our Bayesian analyses. The median provides a more stable estimate of the parameter estimates and is maximally robust against outliers. Additionally, we chose to adopt 90% credible intervals due to suggested convention of Bayesian posterior distributions, as some have speculated that 95% credible intervals can lack stability if insufficient posterior samples are drawn (Kruschke, 2015; McElreath, 2020). It is worth noting that the credible interval reflects the probability that this interval contains the true parameter estimates given our data, or to be more precise, the uncertainty associated with these mediated effects.

#### fMRI Data Acquisition and Preprocessing

The MRI data were acquired on a 3-Tesla Siemens Trio scanner equipped with a 32-channel head coil. A T1-weighed MPRAGE scan was acquired for each participant (TR = 2400 ms, TE =3.16 ms, Flip angle = 8 degrees, 64 slices, slice thickness = 1.0 mm, FOV = 256 x 256 mm). For each EPI bold run (1 or 2 baseline, 6 incentive), 360 volumes were acquired with 4 mm isotropic voxels (TR = 2000 ms, TE = 30.0 ms, Flip angle = 77 degrees, 32 slices, interleaved order, slice thickness=4 mm, FOV = 384×384 mm, acquisition matrix 64×64 yielding an in-plane resolution of 4×4 mm).

Preprocessing for both anatomical and functional data was performed using fMRIPrep version 1.1.7 (Esteban et al., 2019), which is based on Nipype version 1.1.3 (Gorgolewski et al., 2011). The T1-weighted (T1w) image was corrected for intensity non-uniformity (INU) using N4BiasFieldCorrection (ANTs 2.2.0) (Tustison et al., 2010), and used as T1w-reference throughout the workflow. The T1w-reference was then skull-stripped using antsBrainExtraction.sh (ANTs 2.2.0), using OASIS as target template. Brain surfaces were reconstructed using recon-all (FreeSurfer 6.0.1) (Dale et al., 1999), and the brain mask estimated previously was refined with a custom variation of the method to reconcile ANTs-derived and FreeSurfer-derived segmentations of the cortical gray-matter of Mindboggle (Klein et al., 2017). Spatial normalization to the ICBM 152 Nonlinear Asymmetrical template version 2009c (Fonov et al., 2011) was performed through nonlinear registration with antsRegistration (ANTs 2.2.0) (Avants et al., 2008), using brain-extracted versions of both T1w volume and template. Brain tissue segmentation of cerebrospinal fluid (CSF), white-matter (WM) and gray-matter (GM) was performed on the brain-extracted T1w using fast (FSL 5.0.9) (Y. Zhang et al., 2001).

For each of the 8 BOLD EPI runs per participant (across all tasks and sessions), the following preprocessing steps were performed. First, a reference volume and its skull-stripped version were generated using the custom methodology of fMRIPrep. The BOLD reference was then co-registered to the T1w reference using bbregister (FreeSurfer) which implements boundary-based registration (Greve & Fischl, 2009). Co-registration was configured with nine degrees of freedom to account for distortions remaining in the BOLD reference. Head-motion parameters with respect to the BOLD reference (transformation matrices, and six corresponding rotation and translation parameters) were estimated before any spatiotemporal filtering using mcflirt (FSL 5.0.9) (Jenkinson et al., 2002). BOLD runs were slice-time corrected using 3dTshift from AFNI (Cox, 1996, 2012; Cox & Hyde, 1997). The BOLD time-series (including slice-timing correction when applied) were resampled onto their original, native space by applying a single, composite transform to correct for head-motion and susceptibility distortions. These resampled BOLD time-series will be referred to as preprocessed BOLD in original space, or just preprocessed BOLD. The BOLD time-series were resampled to MNI152NLin2009cAsym standard space, generating a preprocessed BOLD run in MNI152NLin2009cAsym space. Several confounding time-series were calculated based on the preprocessed BOLD: framewise displacement (FD), DVARS and three region-wise global signals. FD and DVARS are calculated for each functional run, both using their implementations in Nipype (following the definitions by (Power et al., 2014)). The head-motion estimates calculated in the correction step were also placed within the corresponding confounds file. The BOLD time-series were resampled to surfaces on the following spaces: fsaverage. All resamplings were performed with a single interpolation step by composing all the pertinent transformations (i.e. head-motion transform matrices, susceptibility distortion correction when available, and co-registrations to anatomical and template spaces). Gridded (volumetric) resamplings were performed using antsApplyTransforms (ANTs), configured with Lanczos interpolation to minimize the smoothing effects of other kernels (Lanczos, 1964). Non-gridded (surface) resamplings were performed using mri_vol2surf (FreeSurfer).

The preprocessed BOLD runs were smoothed with a 4 mm FWHM kernel using 3dBlurtoFWHM from AFNI, as well as scaling (voxels were demeaned) using 3dTstat and 3dcalc from AFNI. Finally, the images were reoriented to LPI orientation using AFNI’s 3dresample function.

#### fMRI Data Analysis

To optimize analyses for the mixed block/event-related task design (Petersen & Dubis, 2012; Visscher et al., 2003), a general linear model (GLM) was applied to extract beta estimates and t-statistics for both the sustained liquid block conditions and the transient event-related motivational conditions for each participant. Specifically, AFNI’s 3dDeconvolve function was used to set up the GLM and build an input matrix with the hemodynamic regression model, which contains the duration modulated block effects (dmblock) for baseline, juice, neutral, and saltwater runs, as well as a parameter tent function expansion for every two seconds between 0 and 24 seconds after the onset of the cue stimulus (TENTzero) for each of the 9 motivation conditions (3 levels of money, 3 levels of liquid, rewarded trials only), and unrewarded trials. The TENTzero function eliminates the first and last basis functions from the set, forcing the deconvolved hemodynamic response function (HRF) to be zero at the start and end of the response window. Importantly, this function enables modeling the HRF without assuming a specific shape for the hemodynamic response (e.g., gamma). Such an estimation approach is advantageous for the complex multi-event (i.e., cue, delay, target, feedback) trials employed here.

The GLM was run with 3dREMLfit from AFNI, which performs a generalized least squares time series regression (aka ‘prewhitened’ least squares) combined with a restricted maximum likelihood (REML) estimation of an Autoregressive Moving Average Model (ARMA(1,1)) temporal correlation structure, which has been argued to substantially improve reliability in task fMRI studies (Olszowy et al., 2017). For each participant, we computed whole brain beta estimates for the 4 block types (baseline, juice, neutral, saltwater), 9 tent functions for each of the motivation conditions (3 monetary reward levels x 3 liquid types) (See Equation). A tent function regressor for incorrect trials and the six motion parameters generated from fMRIPrep realignment were included in the GLM as nuisance regressors. Additionally, TRs were censored (current and previous) if the derivative values were estimated (in fMRIPrep) to have a Euclidean norm above .9 mm.

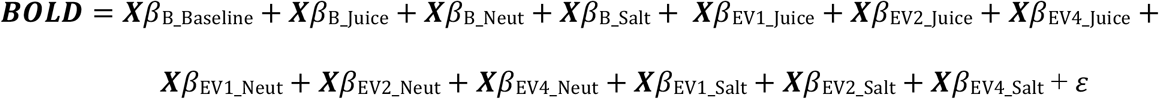

Next, these mixed block/event-related beta estimates were extracted for each of the 400 parcels from the Schaefer cortical atlas, divided into 7 functional networks (Schaefer et al., 2017). The Schaefer atlas was selected because its approach of using a gradient-weighted Markov Random Field to integrate local gradient and global similarity metrics across resting state fMRI and task-based fMRI acquisition protocols yielded the most homogenous parcellations and thus correspond with higher precision to cortical areas compared to prior parcellation schemes (Gordon et al., 2016; Yeo et al., 2011). Additionally, given our a priori hypotheses in subcortical regions, beta estimates were calculated for 19 subcortical regions of interest (ROIs) that were anatomically defined from Freesurfer. Beta estimates were averaged for all voxels within each parcel/region using 3dROIstats from AFNI.

For the dACC ROI analyses, five parcels from the Schaefer cortical parcellation scheme (400 parcels, 7 networks, 2 mm voxels) were identified and averaged, using criteria that were conservative and anatomically focused in corresponding to bilateral dACC based upon a prior meta-analysis on motivated cognitive control (Parro et al., 2018). Specifically, the following parcels were included in bilateral dACC: 107 (‘LH_SalVentAttn_Med_1), 108 (‘LH_SalVentAttn_Med_2’), 110 (‘LH_SalVentAttn_Med_4), 311 (‘RH_SalVentAttn_Med_1’), 312 (‘RH_ SalVentAttn_Med_2’). MNI coordinates of these five parcels corresponding to dACC are listed in the table below.

## Results

### Motivational Incentive Integration Effects: Reward Rate and Self-Reported Motivation Ratings

In the task, participants have the opportunity to earn monetary and liquid incentives on every trial if their performance is accurate and faster than a criterion RT cutoff (defined individually for each participant from their performance in a baseline condition). Consequently, we used the reward rate – defined as the percentage of trials in which the participant earned the available reward – as the primary dependent behavioral measure of motivated cognitive task performance. All participants (n=46) performed above the expected rate of 40%, indicating that they significantly improved their performance relative to baseline levels (45/46 participants showed statistically significant improvements in performance according to a binomial test; successes=117 trials=288, p=.05). Additionally, participants were significantly faster [t(45)=16.298, p<.001] and less accurate [t(45)=7.582, p<.001] during the incentive blocks compared to the baseline block; this shift along the speed-accuracy curve suggests that participants strategically increased their effort in when bundled incentives were added to the task, in order to optimize reward rates. (See Figure 1-2). Finally, RT switch costs (a critical measure of cognitive control) were significantly reduced between baseline and incentive blocks [t(45)=2.956, p=.005], demonstrating overall increased recruitment of cognitive control during incentive blocks relative to baseline blocks. Nevertheless, we focused on reward rate as our primary index of motivated task performance, because it is the most proximal measure indexing the extent to which the expected value of a given trial (i.e., the integrated value of the incentives) could modulate the degree of control specified (as opposed to RT switch costs that typically reflect the preparation of target-related processes and would be most strongly modulated by manipulating the task preparation time (Wylie & Allport, 2000)).

**Figure 1-2:**
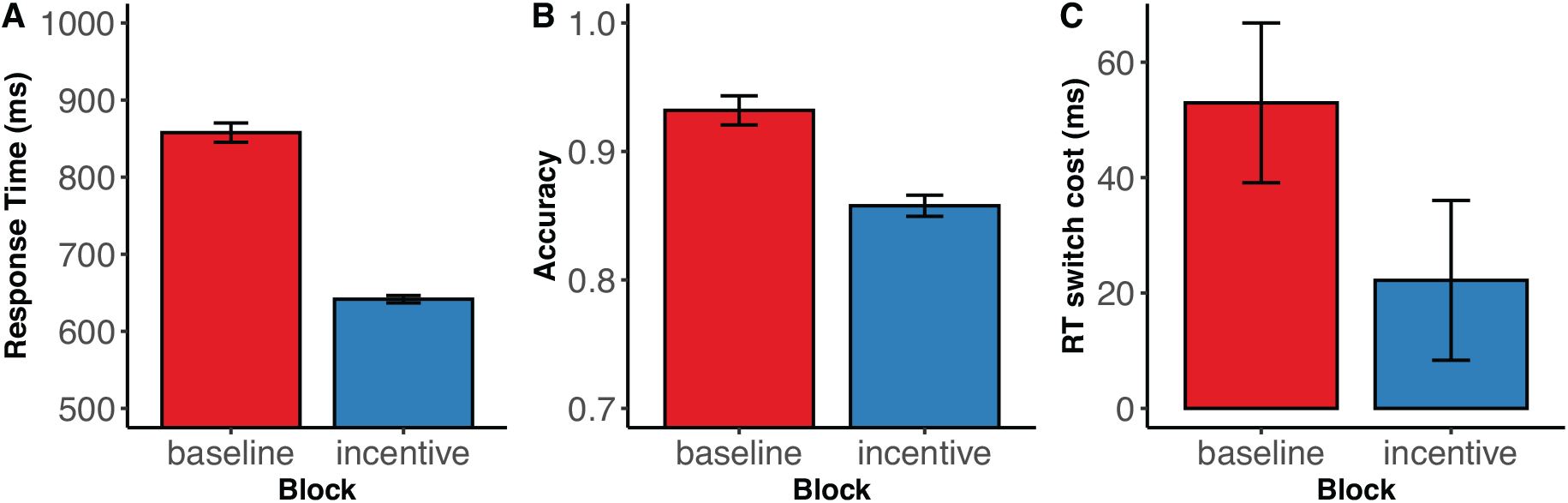
RT, accuracy, and RT switch costs between baseline and incentive blocks. **A)** Comparison of RT in baseline and incentive task blocks demonstrate that motivational incentives are associated with a significant reduction in response times (ms) between baseline and incentive blocks for younger adults [t(45)=16.298, p<.001]. In other words, younger adults are faster with incentives compared to without incentives. **B)** Younger adults showed a significant drop in accuracy between baseline and incentive blocks [t(45)=7.582, p<.001]. Taken together, these data demonstrate that the participants are both faster and more accurate with monetary and liquid motivational incentives. This shift down the speed-accuracy curve to increase reward rate demonstrates that participants increase their effort in accordance with the bundled incentives. All error bars in all plots indicate 95% confidence intervals. **C)** RT switch costs were significantly reduced between baseline and incentive blocks, thus revealing that increased recruitment of cognitive control during the incentive blocks relative to the baseline blocks [t(45)=2.956, p=.005].

**Figure 1-3:**
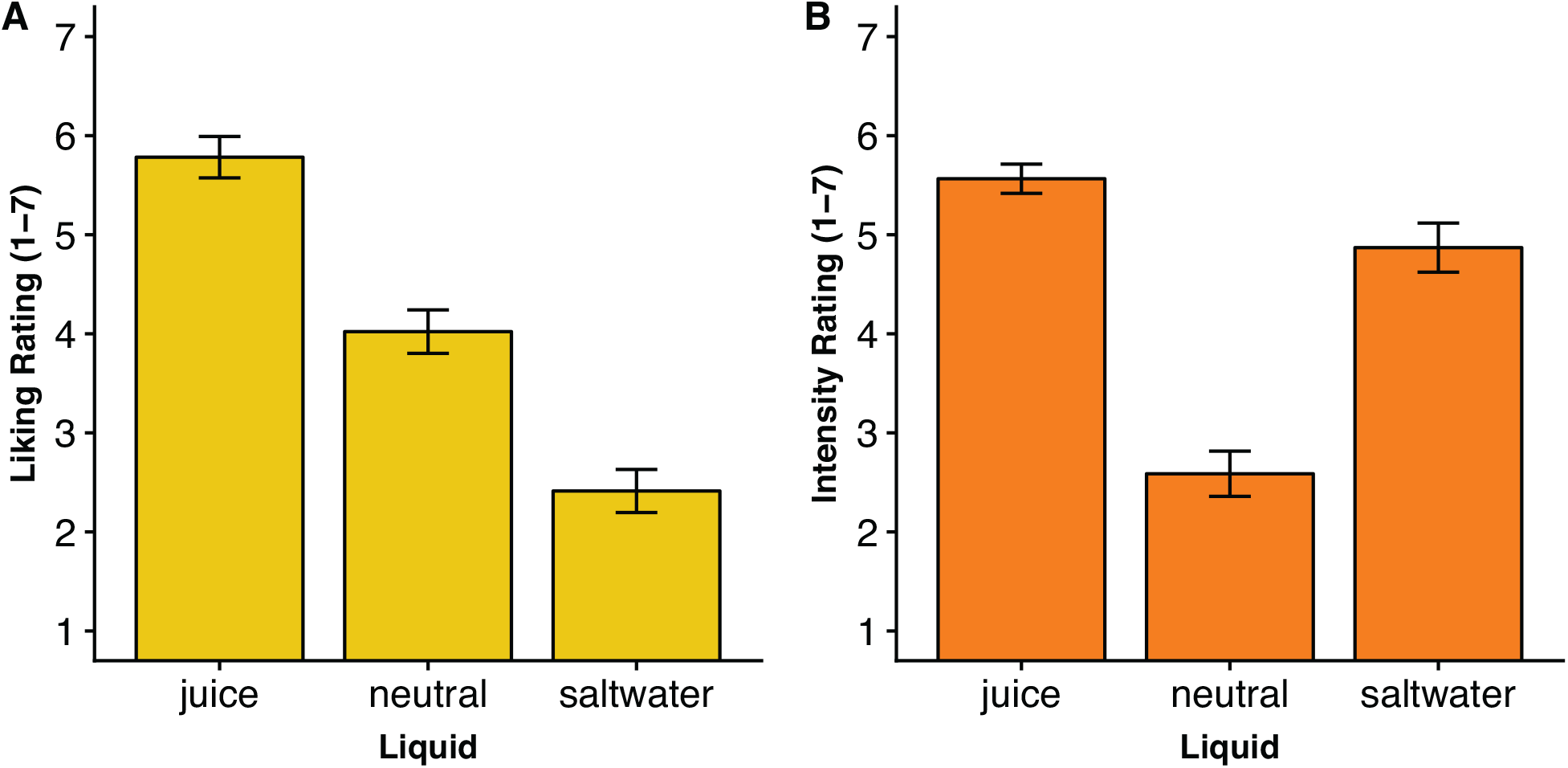
Self-Reported Liking and Intensity Ratings for Each Liquid. **A)** Participants reported significantly different and transitive liquid preferences [b=1.685, t(45)=11.68, p<.001], preferring juice to neutral [t(45)=3.859, p<.001], and neutral over saltwater [t(45)=5.138, p<.001]. B) Participants reported juice and saltwater as more intense than neutral [Juice vs. Neutral: t(45)=12.423, p<.001; Saltwater vs. Neutral: t(45)=6.170, p<.001].

We found that different types of motivational incentives are integrated to modulate cognitive task performance. Specifically, we estimated a general linear mixed model with contrast coded monetary reward (Low = −1, Medium = 0, High = 1) and liquid valence (Saltwater = −1, Neutral = 0, Juice = 1) as fixed effects with participant as a random effect [Reward Rate ~ Money * Liquid + (1 | Subject)]. The model revealed significant effects of both monetary [b=.03, t=4.41, p<.001] and liquid incentives [b=.01, t=2.30, p=.022], but no significant interaction [b=.00, t=−.21, p=.830]. The liquid incentive effects were motivationally valenced, as expected by our prior work, in that performance was better when positive (juice) relative to negative (saltwater) liquids were offered as incentives. Post-hoc analyses further revealed a significant difference between neutral and saltwater [b=.03 t=2.12, p=.034], but no significant effects between juice and neutral [b=.00, t=.08, p=.936]. Thus, the liquid effects on reward rates were primarily driven by impaired performance when saltwater was offered as incentive feedback (Figure 1B; although note that in our prior behavioral studies, we observed both performance facilitation effects due to juice, as well as performance impairments due to saltwater (Crawford et al., 2020; Yee et al., 2016, 2019)). It is noteworthy that we did not detect an interaction between monetary rewards and liquid incentives, suggesting the presence of pure additive effects. Additional analyses of RT and accuracy (using the same linear mixed models as before, except now with RT and accuracy as dependent variables) revealed that reward rate improvements were primarily driven by a faster RT for higher monetary reward [b=-16.25 t=−4.93, p<.001], with no speed-accuracy trade-off (See Figure 1-4 and Table 1-2). However, analyses of switch costs indicated no further effects of monetary rewards and liquid incentives (all p’s > .05, See Table 1-3) beyond the general switch-cost reduction observed in the incentive blocks relative to baseline [RT ~ switch * liquid * money + (1|Subject)]. Switch trials were dummy coded (switch=1, repeat=0), money and liquid were contrast coded same as in previous models. This lack of effect is not particularly surprising, given that switch costs were overall quite small under incentive conditions (~20 ms), and so likely did not have sufficient dynamic range to exhibit sensitivity to the more subtle parametric incentive manipulations. More generally, because of the long cue-target intervals used in the current design, switch costs may not be the most sensitive index of cognitive control for this study, which supports our use of reward rate as the primary indicator of motivated cognitive task performance.

**Figure 1-4:**
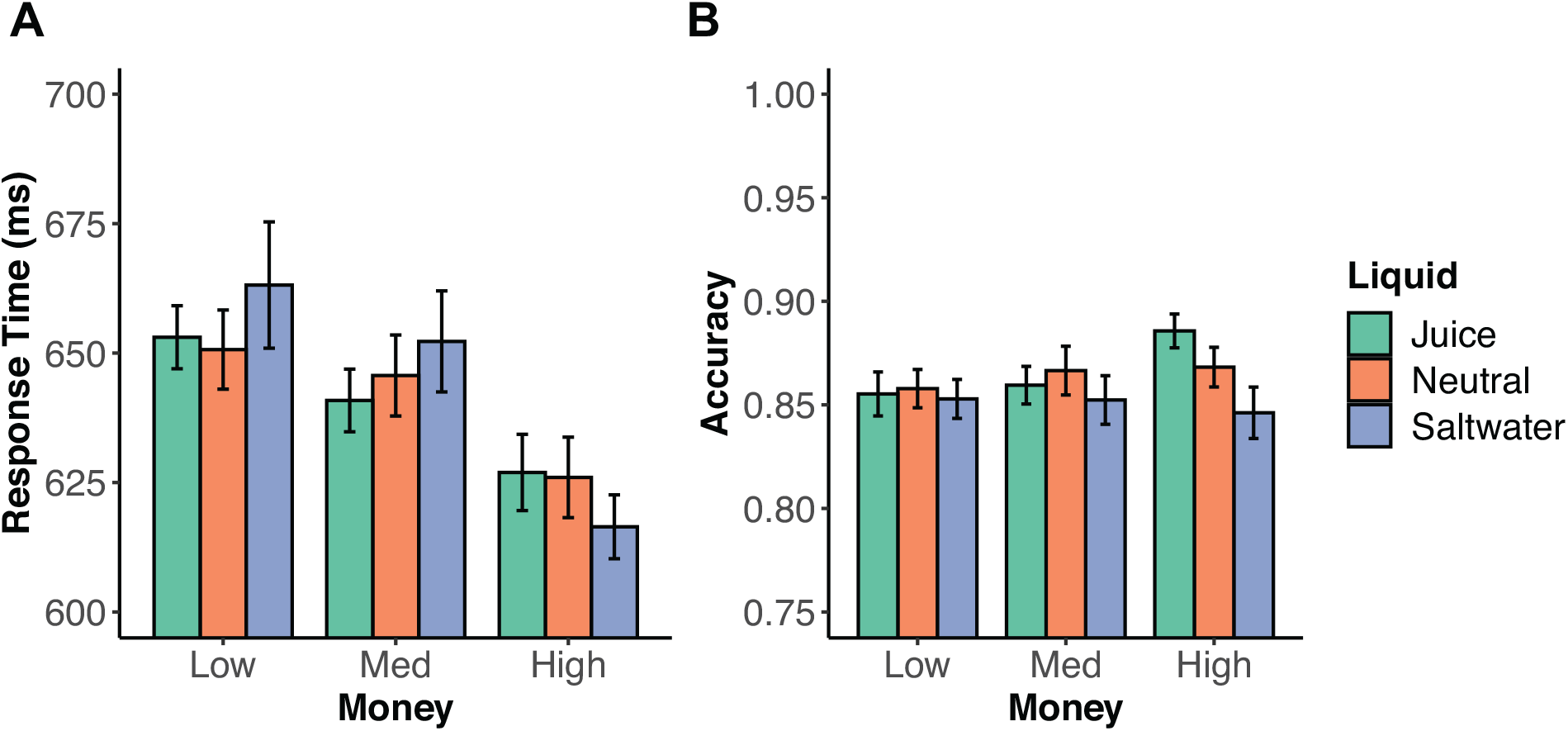
Response Time and Accuracy for each of the 9 Experimental Task Conditions. **A)** Response Time by 9 motivational incentive conditions. Participants were faster on trials with high monetary reward, but there were no differences in liquid incentive, and no significant interaction. **B)** Accuracy by 9 motivational incentive conditions. Accuracy did not differ across monetary reward level, though there was weak effect of liquid incentive that did not meet the threshold for statistical significance. Errors bars indicate 95% confidence intervals.

**Table 1-2:** Response times and accuracy mixed model by monetary reward and liquid incentives. Enhancements in reward rate were primarily driven by faster RT.

| *Predictors* | **Response Time** |  |  | **Accuracy** |  |  |
| --- | --- | --- | --- | --- | --- | --- |
|  | *Estimates* | *Statistic* | *p* | *Estimates* | *Statistic* | *p* |
| Intercept | 641.67 | 34.34 | **<0.001** | 0.86 | 88.02 | **<0.001** |
| Money | -16.25 | -4.93 | **<0.001** | 0.01 | 1.36 | 0.175 |
| Liquid | -1.83 | -0.56 | 0.579 | 0.01 | 1.95 | 0.051 |
| Money:Liquid | 5.14 | 1.27 | 0.202 | 0.01 | 1.81 | 0.071 |
| N | 46<sub>subID</sub> |  |  | 46<sub>subID</sub> |  |  |
| Observations | 414 |  |  | 414 |  |  |
| Marginal R<sup>2</sup> / Conditional R<sup>2</sup> | 0.010 / 0.842 |  |  | 0.012 / 0.450 |  |  |

**Table 1-3:** RT switch cost by monetary and liquid incentives. Notably, we did not observe any incentive effects on switch cots (beyond the general switch-cost reduction observed in incentive blocks relative to baseline in Figure 1–2c). This lack of effect is not surprising, given that switch costs were quite small under incentive conditions (~20 ms), and likely did not have sufficient dynamic range to exhibit sensitive to the more subtle parametric incentive manipulations.

| *Predictors* | **RT (ms)** |  |  |
| --- | --- | --- | --- |
|  | *Estimates* | *CI* | *p* |
| (Intercept) | 629.48 | 592.72 – 666.24 | **<0.001** |
| Switch | 22.56 | 13.67 – 31.45 | **<0.001** |
| Liquid | -5.52 | -13.22 – 2.18 | 0.160 |
| Money | -14.31 | -22.00 – -6.61 | **<0.001** |
| Switch * Liquid | 7.07 | -3.82 – 17.96 | 0.203 |
| Switch * Money | -3.36 | -14.25 – 7.52 | 0.545 |
| Liquid * Money | 0.54 | -8.89 – 9.97 | 0.910 |
| (Switch * Liquid) * Money | 10.23 | -3.10 – 23.57 | 0.133 |
| N<sub>subID</sub> | 46 |  |  |
| Observations | 828 |  |  |
| Marginal R<sup>2</sup> / Conditional R<sup>2</sup> | 0.017 / 0.790 |  |  |

We next examined the relationship between self-report motivation ratings and cognitive task performance. Interestingly, these ratings predicted unique variance in reward rate beyond the experimental manipulation itself. Self-reported measures of motivation were collected for each of the nine trial types (e.g., How motivated were you on Juice $ trials?) after task completion. A linear mixed-effects model [Motivation Ratings ~ Money * Liquid + (1|Subject)] revealed these motivation ratings were significantly predicted by both monetary [b=.69, t=9.31, p<.001] and liquid incentives [b=.75, t=10.04, p<.001], such that participants reported more motivation for higher monetary reward and more appetitive liquid incentives (See Figure 1C). We conducted a hierarchical regression to examine whether self-reported motivation predicted unique variance in reward rate over and above the experimentally manipulated motivational variables [Model 1: Reward Rate ~ Money * Liquid+ (1|Subject); Model 2: Reward Rate ~ Money * Liquid + Motivation Ratings + (1|Subject)]. The hierarchical regression revealed that when self-reported motivation ratings were added to the model with experimental fixed effects (e.g., monetary reward, liquid valence), their inclusion increased predicted reward rate variance over and above the experimentally manipulated motivational variables [χ^2^(3)=20.311, p<.001] (See Table 1-4). These results indicate that self-reported fluctuations in motivation are an important contributor to cognitive task performance in that they predict unique variance beyond the effects of monetary and liquid incentives.

**Table 1-4:**
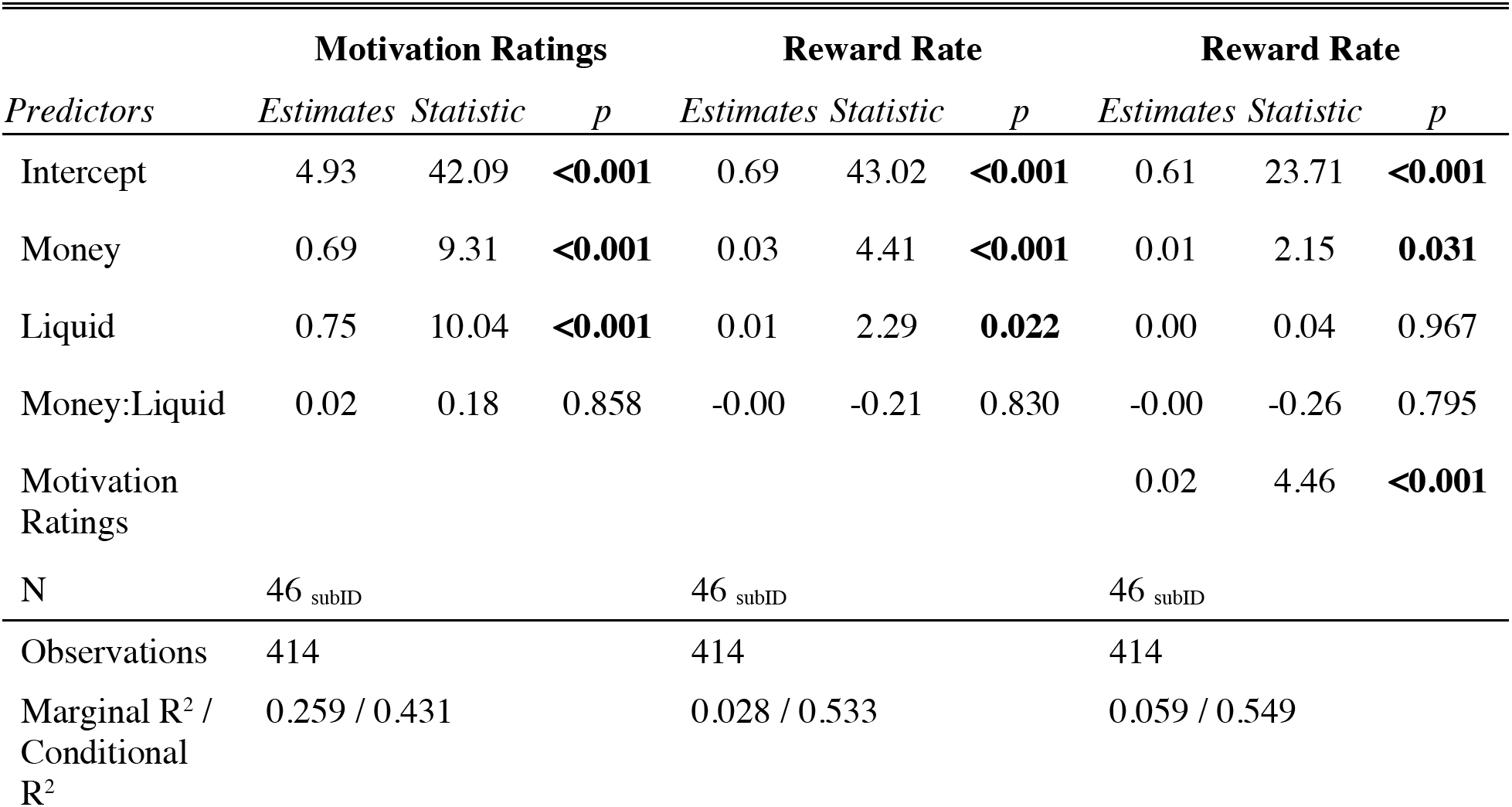
Hierarchical regression with self-report motivation ratings. Motivation ratings were predicted by monetary reward and liquid incentives. Additionally, these motivation ratings significant predicted variance in reward rate in a model including experimental fixed effects (e.g., money, liquid).

Critically, the self-report findings suggest two key interpretations. First, participants could access their subjective motivational states. Second, although these subjective states were modulated by monetary and liquid incentives, the induced motivation from the incentives may have been a more proximal influence on task performance. That is, although both reward rate and self-report motivation ratings are sensitive to the incentive manipulations, the weak association between these metrics [r=.22, t(44)=1.541, p=.13] (See Figure 1D) suggests they might reflect overlapping yet dissociable motivational components of the incentivized cognitive task.

### Dorsal ACC Encodes Both Monetary and Liquid Incentives in Bundled Beta Estimates

We next focused on fMRI data to test whether dACC activity was predicted by monetary and liquid incentives. The dACC ROI encompasses the peak voxels of dACC based upon a prior meta-analysis on motivated cognitive control (Figure 2A), indicating that this ROI is consistently recruited during cognitive control tasks during which a reward incentive can be earned based upon task performance (Parro et al., 2018). For each participant, a general linear model was applied to extract beta estimates for sustained activation for each liquid block condition and event-related activation to the monetary incentive cues for each trial type (estimating activity 4-6 seconds after cue onset), using a deconvolution approach to estimate the time course of activation. Specifically, we averaged the two estimates after the cue onset (t=4s & t=6s) to compute an averaged beta value of the event-related activation associated with each of the monetary incentive cues, accommodating the hemodynamic delay in peak amplitude (Buckner, 1998; Taylor et al., 2018).

As predicted, the results of this analysis revealed that dACC is sensitive to liquid valence via sustained responses and monetary reward via cue-related transient activation during each trial, which is consistent with the incentive delivery structure of the task paradigm and the duration of hemodynamic response lag. To first validate that dACC was independently sensitive to each incentive type, we conducted two linear mixed models with contrast coded monetary reward and liquid valence predicting the block and event-related cue beta estimates, respectively [β_block_~ Money * Liquid + (1 | Subject); β_event_~ Money * Liquid + (1 | Subject)]. The models revealed that the block beta estimates were predicted by liquid incentives [b=.02, t=4.58, p<.001] but not monetary reward [b=0, t=0, p=1.000], whereas the cue beta estimates were predicted by monetary reward [b=.04, t=5.55, p<.001], but not liquid valence [b=0, t=−.31, p=.759]. Next, we “bundled” the beta estimates corresponding to the liquid incentive effects (blocked) and monetary reward effects (event-related, 4-6 seconds after cue onset), which enabled us to calculate nine dACC bundled beta estimates for each participant corresponding to the 9 motivational conditions from the incentive integration task (e.g., $$-Juice, $$$$-Neutral; See Figure 2B). A critical assumption made was that summing the block and event-related beta estimate values would reflect the additive effects of both liquid and monetary incentives in the BOLD signal (i.e., the integrated value). This seems a plausible assumption since the liquid effects were sustained throughout the task block, and thus by definition, available on every trial (i.e., to be added to the cue-triggered event-related activity). Importantly, the mixed model revealed that higher values of dACC bundled beta estimates were associated with higher monetary reward [b=.04, t=4.66, p<.001] and more appetitive liquid incentives [b=.02, t=2.98, p=.003], confirming that neural signals associated with primary and secondary incentives can be combined additively to represent the aggregate motivational value from both incentive types. Paralleling the behavioral results, there was no significant interaction between these two factors [b=0, t=.26, p=.794], suggesting simple additive effects. All model results are shown in extended data Table 1-5.

**Table 1-5:**
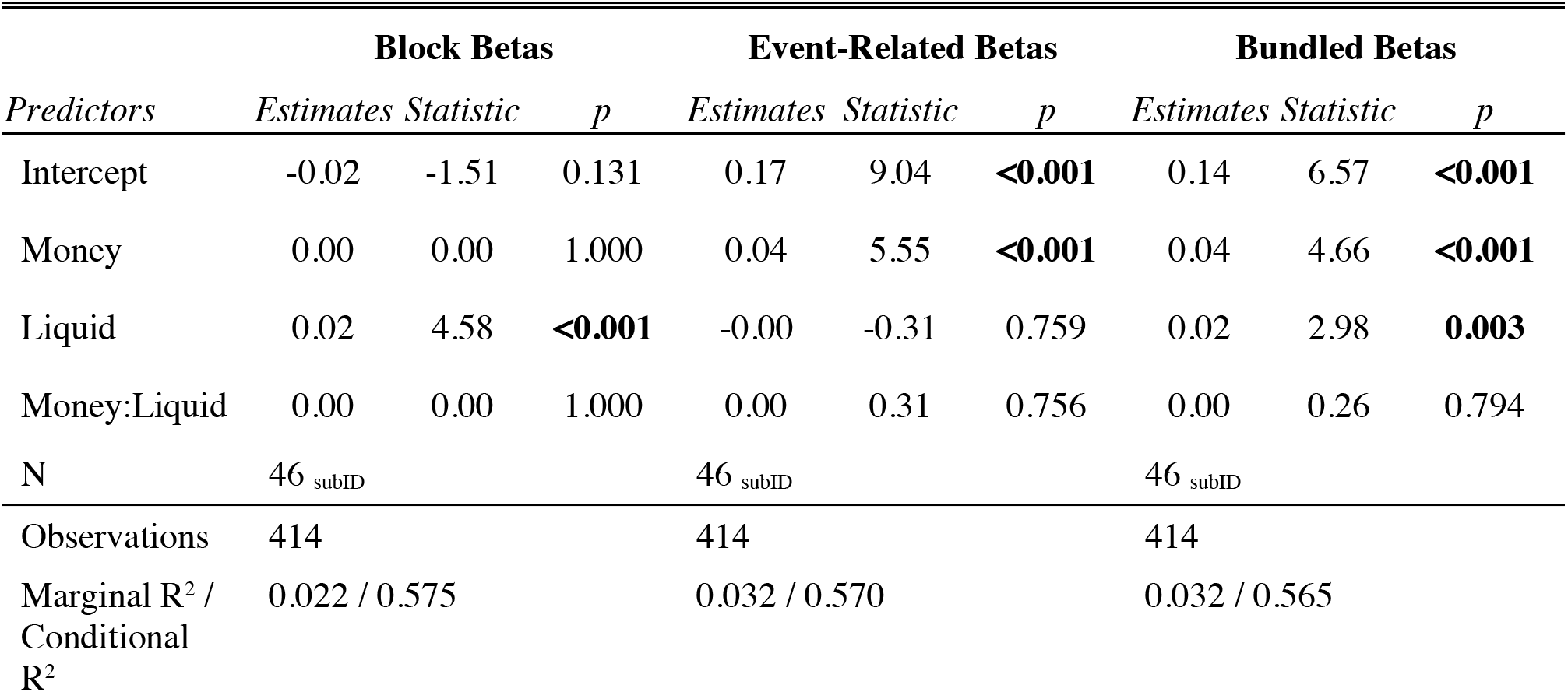
Dorsal Anterior Cingulate Cortex (dACC) beta estimates predicted by monetary reward and liquid incentives.

| *Predictors* | **Block Betas** |  |  | **Event-Related Betas** |  |  | **Bundled Betas** |  |  |
| --- | --- | --- | --- | --- | --- | --- | --- | --- | --- |
|  | *Estimates* | *Statistic* | *p* | *Estimates* | *Statistic* | *p* | *Estimates* | *Statistic* | *p* |
| Intercept | -0.02 | -1.51 | 0.131 | 0.17 | 9.04 | **<0.001** | 0.14 | 6.57 | **<0.001** |
| Money | 0.00 | 0.00 | 1.000 | 0.04 | 5.55 | **<0.001** | 0.04 | 4.66 | **<0.001** |
| Liquid | 0.02 | 4.58 | **<0.001** | -0.00 | -0.31 | 0.759 | 0.02 | 2.98 | **0.003** |
| Money:Liquid | 0.00 | 0.00 | 1.000 | 0.00 | 0.31 | 0.756 | 0.00 | 0.26 | 0.794 |
| N | 46<sub>subID</sub> |  |  | 46<sub>subID</sub> |  |  | 46<sub>subID</sub> |  |  |
| Observations | 414 |  |  | 414 |  |  | 414 |  |  |
| Marginal R<sup>2</sup> / Conditional R<sup>2</sup> | 0.022 / 0.575 |  |  | 0.032 / 0.570 |  |  | 0.032 / 0.565 |  |  |

### Reward Rate Predicted by Dorsal ACC Bundled Betas

Given the hypothesized role of dACC as a region that integrates various sources of motivational value into a modulatory signal that adaptively allocates cognitive control (Shenhav et al., 2013, 2017), we aimed to test whether experimentally-manipulated within-subjects variability in dACC was significantly associated with similar within-subject variability in reward rate, our metric of motivated cognitive task performance. Such an association would provide compelling evidence demonstrating that dACC computes the aggregate value of both motivational incentives and predicts motivated task performance. We conducted a linear mixed effects model with reward rate predicted by dACC bundled beta estimates and ‘incentive conditions’ (contrast coded as the sum of liquid and monetary reward contrast codes). Our analyses revealed that reward rate was significantly predicted by both incentive conditions (as expected from prior analyses) [b=.02, t=2.91, p<.001], but also the dACC bundled betas [b=.09, t=2.41, p=.016]. A critical and important observation is that the inclusion of dACC bundled betas explained significant additional variation in reward rate beyond the experimentally-manipulated incentive effects, suggesting a role of dACC in encoding subjective motivational value. Moreover, these data are consistent with the EVC account of dACC as a hub for integrating value and cost information to derive an optimal control signal that balances task demands against potential rewards.

### Self-Report Motivation Ratings Predicted by Dorsal ACC Bundled Betas

A secondary aim was to understand the putative relationship between the extent to which motivated cognitive task performance (i.e., reward rate) and self-reported motivation measures may be encoded in dACC activation. As self-report measures of motivation have been historically viewed as the “gold standard” for assessing human motivational states (Hermans, 1970), we investigated the extent to which human dACC neural signals may encode self-reported motivation to exert cognitive control in order to maximize earnings of the bundled incentives. Here, we conducted a similar mixed model as above except now with self-report motivation ratings (rather than reward rate) as the dependent measure. Our analyses revealed that self-reported motivation ratings were predicted by both incentive condition [b=.67, t=9.78, p<.001] and dACC bundled betas [b=1.64, t=3.96, p<.001]. These results are particularly intriguing because this association reveals that dACC neural signals also reflect a subjective (and accurate) measure of the valuation of the bundled motivational incentives in the current task context. Moreover, such ratings may tap into a complementary motivation-related construct distinct from motivation associated with the exerted effort on a cognitively demanding task.

### Dorsal ACC Effects on Reward Rate Mediated by Self-Report Motivation Ratings

Given the observed strong associations between dACC and reward rate, as well as between dACC and self-reported motivation ratings, we next examined the putative relationship between within-subjects variability across different incentive conditions in dACC, reward rate, and motivation ratings. In particular, given the utility of self-report ratings as a powerful metric for measuring subjective motivational states, we hypothesized that such ratings might partially explain the proximal motivational impact of the incentives on the associative modulatory relationship between dACC activity and reward rate. Such a within-subjects mediation would be compelling, as it would reveal the extent to which dACC may encode these two behavioral metrics as similar or distinct indices of an individual’s current motivational state.

First, we conducted three linear mixed models and confirmed that experimentally manipulated incentives demonstrated quantifiable effects on within-subjects variability in dACC, reward rate, and self-report ratings. As indicated previously, incentive conditions significantly predicted dACC bundled betas [b=.03, t=4.04, p<.001], reward rate [b=.02, t=3.36, p<.001], and motivation ratings [b=.72, t=10.35, p<.001]. (See red dashed lines in Figure 3). Thus, these data demonstrate that the incentive manipulations influence all three variables, while also suggesting that these motivational variables may be mediated or moderated by each other.

**Figure 3:**
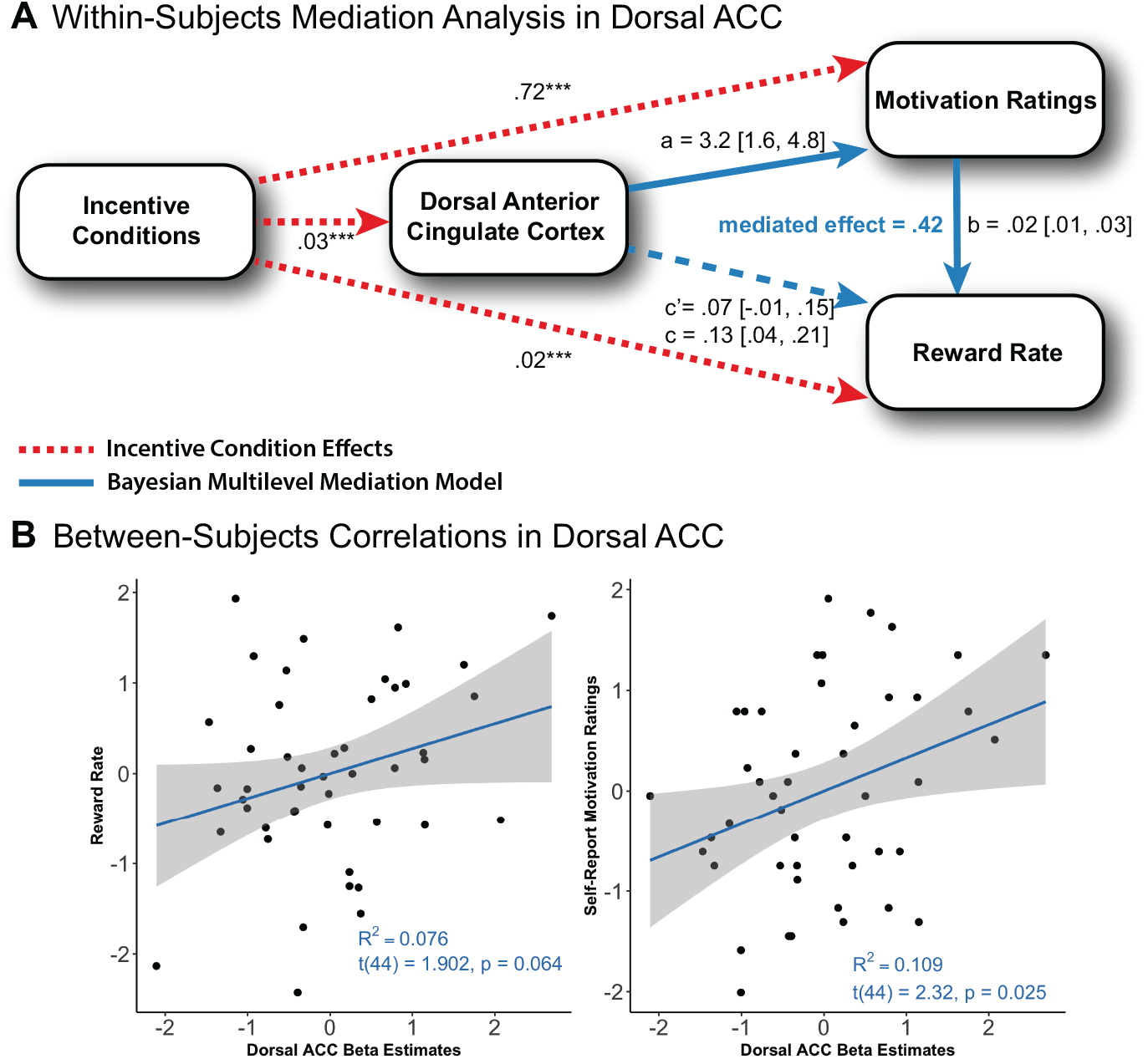
Dorsal ACC Effects on Reward Rate Mediated by Self-Report Motivation Ratings. **A)** Within-Subjects Mediation Analysis of Dorsal ACC. Three linear mixed effects models were conducted to confirm the effects of incentive conditions in dACC, reward rate, and motivation ratings (See red dashed lines). The models revealed that incentive conditions significantly predicted dACC bundled betas, reward rate, and motivation ratings. Next, a Bayesian multilevel within-subjects mediation analysis was conducted to test for a relationship between these three variables. The mediation analyses revealed that the relationship between within-subjects variability in dACC and reward rate was mediated by within-subjects variability in motivation ratings (See blue lines). Inclusion of motivation ratings partially mediated the association between dACC and reward rate proportion mediated effect=.42, CI_90_=(.06,1.10). 90% credible intervals are used as the range of the posterior distribution for each of the path parameters. *p<.05; **p<.01; ***p<.001. **B)** Between-Subjects Correlations of Dorsal ACC. The left scatterplot of z-scored averaged dACC beta estimates and reward rate by subject revealed an association between dACC and reward rate (though beneath threshold for statistical significance). The right scatterplot shows z-scored averaged dACC beta estimates and self-report motivation ratings also revealed association between dACC and motivation ratings.

Next, we conducted a Bayesian multilevel mediation analysis and observed that incentive-modulated effects on the relationship between dACC and reward rate were mediated by such variability in motivation ratings, as prior linear mixed models confirmed incentive effects across all three variables. The within-subjects mediation analysis utilized averaged values across each of the nine possible motivational incentive conditions for each participant. This enabled us to compare within-subjects variability across incentive conditions between these variables (dACC, reward rate, motivation ratings). Notably, this approach eliminated the need for including contrast-coded incentive effects in linear mixed models that were used to validate assumptions about associations between these variables prior to conducting the mediation analysis. Linear mixed models (without contrast-coded incentive effects) confirmed that dACC activity significantly predicted both reward rate [b=.14, t=3.77, p<.001] and motivation ratings [b=2.77, t=5.98, p<.001]. Moreover, in a linear mixed model of reward rate with motivation ratings as the mediator, controlling for the predictor revealed that motivation ratings significantly predicted reward rate [b=.02, t=5.53, p<.001], and more importantly, the inclusion of these ratings weakened the effect of dACC, though it still remained statistically significant [b=.08, t=2.15, p=.032].

These variables were submitted to a Bayesian multilevel mediation (using the bmlm package in R) to estimate the medians and 90% credible intervals for the posterior distributions for each path parameter (See blue lines in Figure 3). The mediation analysis revealed that motivation ratings partially mediated the relationship between dACC and reward rate [mediated effect=.05, CI_90_=(.01,.10); proportion mediated effect=.42, CI_90_=(.06,1.10)], significantly reducing the direct effect [c=.13, CI_90_=(.04,.21); c’=.07, CI_90_=(-.01,.15)]. We also conducted an alternative model which revealed a weaker partially mediated effect of reward rate on the relationship between dACC and motivation ratings [mediated effect=.44, CI_90_=(.06,.90); proportion mediated effect=.13, CI_90_=(.02,.31)], which, critically, did not significantly reduce the direct effect [c=3.30, CI_90_=(1.73,4.85); c’=2.84, CI_90_=(1.32,4.34)]. Thus, although dACC is significantly associated with incentive motivational task performance, this relationship is partially mediated by the variance explained by the self-report motivation ratings. Notably, these mediation models reveal that the subjective motivational incentive value is the more proximal factor that modulates task performance, rather than the incentive-driven performance modulation driving the motivation ratings. As such, these results provide compelling evidence that dACC encodes the integrated incentives primarily in terms of their subjective motivational value, which in turn modulates task performance. Parameter estimates are listed in Table 1-5 and Table 1-6.

**Table 1-6:** Estimated parameters from Bayesian multilevel mediation analysis of motivation ratings mediating relationship between dACC and reward rate.

| **Parameter** | **Mean** | **SE** | **Median** | **5%** | **95%** | **n_eff** | **Rhat** |
| --- | --- | --- | --- | --- | --- | --- | --- |
| **a** | 3.24 | 0.97 | 3.24 | 1.62 | 4.83 | 18930 | 1 |
| **b** | 0.02 | 0.00 | 0.02 | 0.01 | 0.03 | 17809 | 1 |
| **cp** | 0.07 | 0.05 | 0.07 | -0.01 | 0.15 | 16460 | 1 |
| **me** | 0.05 | 0.03 | 0.05 | 0.01 | 0.10 | 7949 | 1 |
| **c** | 0.12 | 0.05 | 0.13 | 0.04 | 0.21 | 22171 | 1 |
| **pme** | 0.54 | 7.17 | 0.42 | 0.07 | 1.13 | 19967 | 1 |

In addition to examining within-subjects variability, we also tested whether between-subjects’ differences in dACC activation were associated with reward rate and motivation ratings. That is, if baseline differences in overall dACC activation across individuals correlated with reward rate and motivation ratings, it would suggest that dACC also reflects more stable levels of task engagement (between-subjects variability), beyond integrating task-related changes in incentive values towards motivated task performance (within-subjects variability). We averaged and normalized subject-level values for dACC activation, reward rate, and motivation ratings (i.e., rather than across the 9 incentive conditions). Our results revealed a moderate association between dACC and reward rate [r=.28, t(44)=1.902, p=.064], as well as between dACC and motivation ratings [r=.33, t(44)=2.32, p=.025] (See Figure 3B). Individuals with higher average dACC activity showed a trend towards reporting overall higher motivation ratings (and to a lesser extent, achieved higher reward rates).

These between-subjects dACC effects were not as reliable as the within-subjects dACC effects previously demonstrated. In particular, when both subject-level dACC activity and motivation ratings were included in a model predicting subject-level individual differences in average reward rate, neither variable was statistically reliable [dACC: b=.23, t=1.47, p=.149; Motivation Ratings: b=.15, t=.99, p=.328]. Thus, our data suggest that the dACC accurately encodes context-specific motivational states that are primarily modulated in a dynamic fashion (i.e., by incentive cues) rather than by more stable or global trait-like differences in motivation. Notably, although the utilization of an *a priori* ROI as well as hypothesized motivational processes, reduced potential false positive concerns regarding the between-subjects effects, these results should still be interpreted with caution. The modest sample size of the current study (N=46) suggests that it is potentially underpowered to detect individual differences effects in dACC, or alternatively, could produce a biased inflation of the estimated effects (Cremers et al., 2017; Dubois & Adolphs, 2016). Validation of the individual differences’ relationships between dACC and motivated processes would require collecting a sufficiently powered sample (Barch et al., 2013; Turner et al., 2018) and/or conducting *a priori* power analyses with given effect sizes to determine the estimated reliability of the findings expected in a novel sample (Mumford, 2012).

### Dorsal ACC Selectively Encodes Subjective Motivational Value and Modulates Motivated Task Performance

Finally, we complemented our analyses of dACC activity patterns by examining other relevant brain regions of interest. In particular, a number of well-established brain regions have been implicated in value-based decision making (e.g., striatum, vmPFC), and other reward-related processes, such as taste processing (e.g., anterior insula). Consequently, we tested whether the subjective motivational value signal associated with task performance was also encoded in these other value-related brain regions. We conducted the same linear mixed models with reward rate and motivation ratings predicted by the motivational incentive conditions and bundled betas, except now using bundled betas from other regions of interest associated with value-based decision making (Bartra et al., 2013; Sescousse et al., 2013, 2015) as well as taste processing (Small, 2010). Of these selected ROIs, it is noteworthy that only dACC significantly predicted reward rate. In contrast, motivation ratings were additionally significantly predicted by caudate [b=.74, t=2.29, p=.022], putamen [b=1.29, t=3.22, p=.001], and anterior insula [b=1.36, t=2.87, p=.003] (See Figure 4). These data reveal that although several regions appear to predict motivation ratings (suggesting multiple brains regions may track the subjective motivation of the incentives), the specificity of the association between dACC and reward rate robustly supports the EVC theory, i.e., that dACC may play a significant and selective role in modulating how motivational values are translated in effortful action during the performance of a cognitively demanding task.

**Figure 4:**
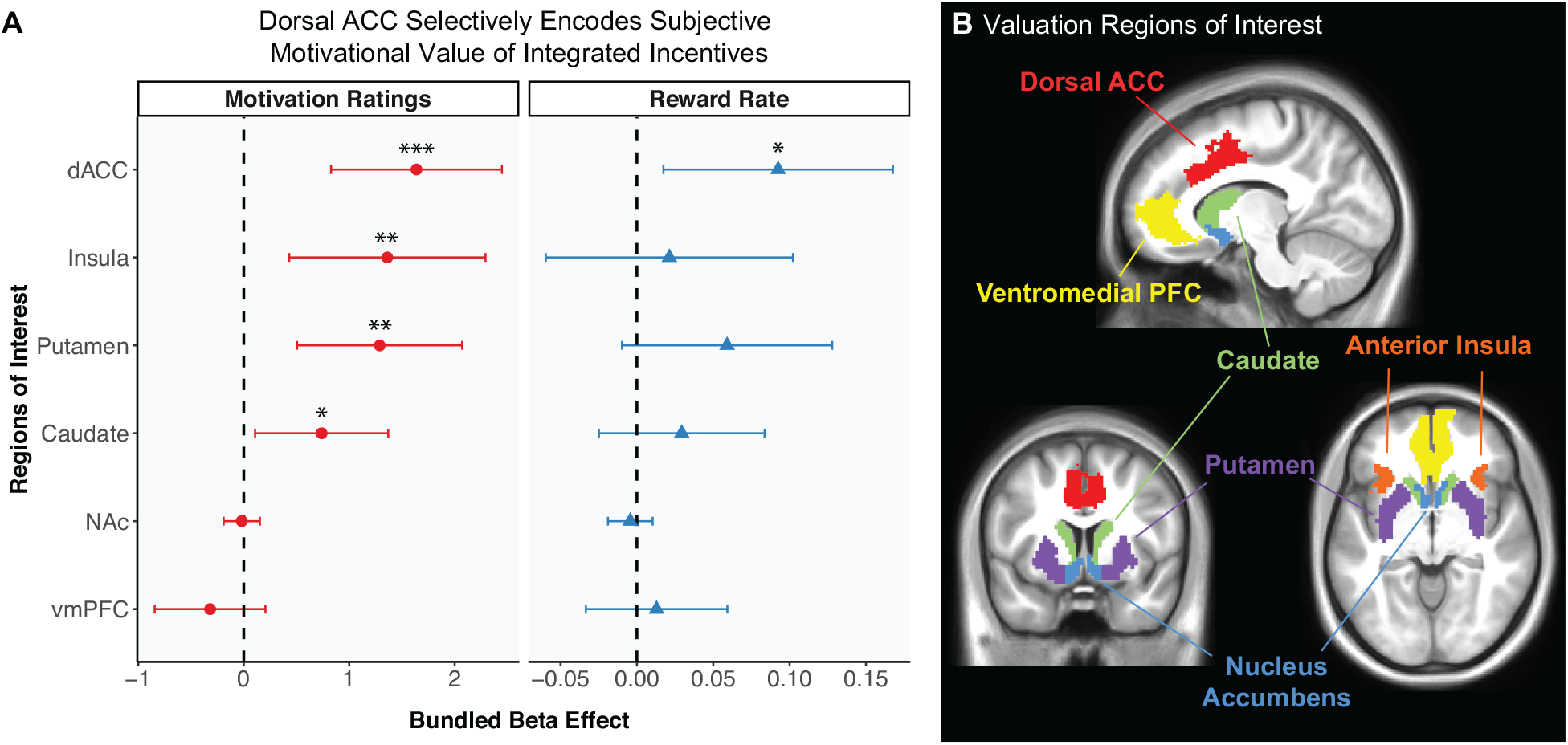
Dorsal ACC Selectively Encodes Subjective Motivational Value and Modulates Motivated Task Performance. **A)** To test for whether motivational incentive integration effects were present in other brain regions, bundled beta estimates were calculated for *a priori* regions of interest associated with value-based decision-making (striatum, vmPFC) and taste processing (anterior insula). Mixed models were implemented with selected ROIs and plotted to compared to dACC effects on reward rate and motivation ratings. Error bars represent 95% confidence intervals. The right panel reveals that only dACC significantly predicted reward rate. The left panel reveals that motivation ratings were also significantly predicted anterior insula, caudate, and putamen, in addition to dACC, suggesting that self-reported motivation of incentives is encoded across multiple valuation brain regions. These data reveal the specificity of the relationship between dACC and reward rate, demonstrating that dACC encodes the translation of motivational value is transformed into effortful actions on a cognitively demanding task. *p<.05; **p<.01; ***p<.001. **B)** Visualization of valuation brain regions of interest.

## Discussion

We found novel evidence that dACC represents the integrated subjective motivational value of bundled primary and secondary incentives, and moreover, that this bundled neural signal is associated with changes in motivated cognitive task performance. We leveraged key features of our incentive integration paradigm, which enables estimation of the combined effect of both monetary and liquid incentives on brain activity and task performance, while also allowing for examination of motivational valence effects through the use of both appetitive and aversive liquid feedback. Our findings support predictions of the Expected Value of Control (EVC) framework (Shenhav et al., 2017), which postulates that dACC integrates the motivational value of potential outcomes (e.g., the appetitive value of earning monetary reward subtracting the aversive value of the saltwater) to determine the optimal allocation of cognitive control. We find clear evidence that dACC acts as an integrative motivation-cognition hub, thus facilitating the pursuit of cognitively effortful goals (Holroyd & Yeung, 2012). Although prior work has demonstrated the role of dACC in utilizing reward information to optimally allocate effort in rodents (Holroyd & McClure, 2015; Hosking et al., 2014), ours is the first study to demonstrate the role of human dACC in encoding subjective motivational value across integrated incentives to modulate behavioral performance in a task with high cognitive demands. In particular, consistent with EVC predictions, we found that reward rate tracked the subjective motivational value of cognitive control over task performance (i.e., in terms of both the integrated incentives available and self-reported motivation), and that this relationship was mediated by fluctuations in dACC activity.

A notable observation was the discovery that activity in the dACC was associated with not only task performance, but also with self-reported motivation, which is intriguing since these motivation ratings explained unique variance in reward rate beyond the incentive manipulations. Additionally, these motivation ratings partially mediated the relationship between dACC and reward rate, providing suggestive evidence that such ratings may reflect the more proximal impact of the incentive conditions encoded in dACC activity. Moreover, the self-reported motivational effects of the incentive manipulations may be somewhat distinct, though functionally related to the motivational effects inferred from behavioral enhancements observed during incentivized task performance (Bonner and Sprinkle, 2002). Alternatively, these motivation ratings may simply reflect a more accurate subject-specific indication of incentive salience based upon the ordinal levels from the manipulation.

It is important to acknowledge a limitation regarding the inclusion of self-report motivation ratings. Although the use of these ratings was prompted by our group’s prior work demonstrating that they predict unique variance beyond experimentally manipulated incentive effects (Yee et al., 2016, 2019), they do not capture the dynamic variability likely present in subjective motivation, as it gets modulated by performance, feedback, or physiological states (e.g., satiety). Although some have examined the dynamic variability in self-reported mood and behavior over the span of days/weeks (Moskowitz & Young, 2006; Rutledge et al., 2014), how subjective motivation varies dynamically throughout the course of a behavioral task context remains to be explored. Future work should aim to investigate the extent to which both measures (i.e., motivated task performance, self-report) reflect convergent versus divergent neural signals underpinning motivation, and explore the temporal dynamics of motivation within a task. It is important to note that including both measures simultaneously incurs its own set of limitations, and would require consideration of how self-report probes potentially alter the demand characteristics of motivated task performance (Polivy & Doyle, 1980; Velten, 1968). Nevertheless, such investigations have the potential to reconcile the frequently observed discrepancies between how both measures (self-report, behavioral performance) reflect motivated cognition (Dang, King and Inzlicht, 2020). As such, this work could advance current understanding of how distinct sources of motivational information are represented in the brain.

Interestingly, we found evidence of an incentive integration neural mechanism present in the dACC, but not in other brain regions classically associated with representing the neural common currency of subjective value (Levy & Glimcher, 2012; Peters & Buchel, 2010). It is possible that these other regions may also be involved in integrating incentive value, but that the effects were not detected in the ROI-based analysis employed here, and could require methods with greater granularity to detect putative heterogeneous processes within these ROIs. Nevertheless, our results are consistent with prior work showing a distinction between the subjective value signals associated with explicit economic choices (i.e., choosing one good over another) versus when such values are relevant for behavioral actions (Cai & Padoa-Schioppa, 2012; Camille et al., 2011; Kolling et al., 2016). The former process (value-based decision-making) involves explicit valuation and comparison of incentives, whereas the latter process (motivated cognitive control) involves implicit processing and utilization of incentive value for adaptive mobilization of cognitive control. Although a key feature of this study is the utilization of both monetary and liquid incentives, our task significantly deviates from prior studies evaluating the expectation of primary and secondary incentives (Chib et al., 2009; Kim et al., 2011; S. Q. Park et al., 2012; Z. Zhang et al., 2017).

Most prior evidence supporting the neural common currency account arises from human and animal studies which include an explicit valuation phrase or when an economic choice is required between available goods (Fromer et al., 2019; Padoa-Schioppa & Conen, 2017). Because our task paradigm is optimized for motivated cognitive control, it is unsurprising that only dACC appeared to encode the incentive condition effects on reward rate. These results are consistent with prior work demonstrating that cost-benefit valuation in physical effort tasks elicit activation in dACC, but not vmPFC (Croxson et al., 2009; Klein-Flugge et al., 2016). Furthermore, this distinction is supported by our observation that motivation ratings were associated with bundled betas in dorsal striatum (e.g., caudate, putamen), a region well-known to be associated with motivated action selection (Balleine et al., 2007; Balleine & O’Doherty, 2010; Miller et al., 2014). Future work could bridge this gap via including an additional valuation phase for bundled incentives (e.g., $$$$ reward + saltwater) or a choice component, in which preferences can be expressed (e.g., $$$$ reward + saltwater vs. $$ reward + juice). Such a paradigm might more clearly reveal regions involved in value-based decision-making (e.g., vmPFC, ventral striatum) and/or motivated cognitive control (e.g., dACC, dorsal striatum).

A broader question relates to developing a mechanistic understanding of how dACC integrates incentives during motivated cognitive control. The dACC contains a heterogeneous population of neurons and underpins a diverse array of cognitive, motor, and affective functions (Bush et al., 2002; Heilbronner & Hayden, 2016; Vega et al., 2016). However, in light of our key findings supporting the role of dACC as a hub for motivation-cognition interactions by integrating the combined appetitive and aversive values of diverse incentives (Parro et al., 2018), the precise calculations by which different neurons, voxels, or sub-regions within the dACC perform incentive value integration remains to be elucidated. One possibility is that distinct neural patterns or voxel clusters within dACC may distinctly encode positive outcomes (e.g., monetary rewards, juice) and negative outcomes (e.g., punishments, saltwater). Such patterns seem plausible, given recent work demonstrating that dACC neurons respond to rewards and punishments in nonhuman primates (Monosov, 2017; Monosov et al., 2020) and rodents (Schneider et al., 2020). Alternatively, dACC voxels may be multiplexed to encode both positive and negative outcomes, or even context-specific incentives and actions (Hayden & Platt, 2010). An important future direction would be investigation with higher spatial resolution of when and how dACC integrates diverse incentives to represent the subjective motivational value in cognitive control contexts.

Finally, developing greater insight into how incentive motivation is integrated and represented provides a crucial foundation from which to elucidate the neural mechanisms underpinning how subjective value signals modulate cognitive task representations in prefrontal cortex (PFC). Whereas prior work has shown that lateral PFC signals are enhanced when higher monetary rewards are present (Bahlmann et al., 2015; Dixon & Christoff, 2012; Duverne & Koechlin, 2017; Kouneiher et al., 2009), how such cognitive task PFC representations interface with the subjective motivational value computation in dACC remains an open question. Multivariate analyses (e.g., decoding or representational similarity analysis) could be exploited to probe how value signals are integrated (in dACC) to modulate neural representations of cognitive task rules (in lateral PFC) and ultimately translated to measurable changes in behavior (Etzel et al., 2015; Wisniewski et al., 2015). Such findings could inform systems-level neural representations of how diverse motivational incentives combine into subjective motivational value to support cognitive task goals.

Broadly, given the diversity of incentives that people regularly encounter (e.g., social rewards), an open question relates to understanding how people evaluate and integrate multiple diverse incentives in the real world (Lehner et al., 2017). Our innovative approach provides an initial step towards more careful study of real-world effort allocation, through which incentives can vary along both categories (e.g., monetary, liquid, social) and dimensions (e.g., appetitive, aversive), and are seamlessly integrated (e.g., consideration of both monetary rewards and social praise (Crawford et al., 2020; H. R. P. Park et al., 2018)). Importantly, such insight into the neural mechanisms underpinning how motivational incentives are integrated to modulate effortful tasks can provide a mechanistic framework to advance understanding of how motivational or cognitive deficits arise in clinical disorders, such as anxiety, depression (Clery-Melin et al., 2011; Grahek et al., 2019; Huang et al., 2015) or addiction (Koob & Moal, 2008; Volkow et al., 2017).

## Acknowledgments

The research was supported by the National Institutes Health (R21-AG058206, R21-AG067295 to T.S.B. and subaward R24-AG054355 D.M.Y.) and McDonnell Center for Systems Neuroscience (T.S.B. and D.M.Y.). D.M.Y. and J.L.C. were supported by T32-AG000030, and

D.M.Y. was additionally supported by F31-DA042574 and T32-NS073547. We would like to thank Carolyn Dean Wolf, and Katherine Shapiro for their assistance with data collection and technical support. We also thank Amitai Shenhav, Katherine Conen, as well as the CCP and Shenhav labs for helpful input and discussions during manuscript preparation.

---

**Table 1:** Parcels included in Bilateral Dorsal ACC ROI Mask.

| Parcel ID | Parcel Number | MNI Coordinates (RAS) |  |  |
| --- | --- | --- | --- | --- |
|  |  | x | y | z |
| **107** | LH_SalVentAttn_Med_1 | -6 | 22 | 32 |
| **108** | LH_SalVentAttn_Med_2 | -6 | 0 | 40 |
| **110** | LH_SalVentAttn_Med_4 | -6 | 10 | 48 |
| **311** | RH_SalVentAttn_Med_1 | 8 | 18 | 26 |
| **312** | RH_SalVentAttn_Med_2 | 8 | 2 | 42 |

**Table 1-7:** Estimated parameters from Bayesian multilevel mediation analysis of reward rate mediating the relationship between dACC and motivation ratings.

| **Parameter** | **Mean** | **SE** | **Median** | **5%** | **95%** | **n_eff** | **Rhat** |
| --- | --- | --- | --- | --- | --- | --- | --- |
| **a** | 0.12 | 0.05 | 0.12 | 0.04 | 0.21 | 19517 | 1 |
| **b** | 3.46 | 0.76 | 3.46 | 2.22 | 4.72 | 20403 | 1 |
| **cp** | 2.83 | 0.92 | 2.84 | 1.32 | 4.34 | 11036 | 1 |
| **me** | 0.45 | 0.26 | 0.44 | 0.06 | 0.90 | 13288 | 1 |
| **c** | 3.29 | 0.95 | 3.30 | 1.73 | 4.85 | 11467 | 1 |
| **pme** | 0.15 | 0.16 | 0.13 | 0.02 | 0.31 | 18185 | 1 |

